# Stimulus predictability rather than inter-trial interval itself affects motor evoked potential amplitudes

**DOI:** 10.64898/2026.09.12.750930

**Authors:** Saman Seifpour, Nele Haferkorn, Suhas Vijayakumar, Til Ole Bergmann

## Abstract

The inter-trial interval (ITI) of transcranial magnetic stimulation (TMS) can modulate motor-evoked potential (MEP) amplitude and may thus confound assessments of corticospinal excitability. We tested whether this effect depends on the ITI duration per se or the associated temporal predictability and whether intracortical inhibitory processes mediate these effects on. In 26 healthy participants, we recorded single-pulse MEPs and paired-pulse measures of short-interval intracortical inhibition (SICI; 2-ms interstimulus interval) and long-interval intracortical inhibition (LICI; 100-ms interstimulus interval) at ITIs of 2, 5, and 10 s. ITIs were presented in different Patterns, i.e., separate fixed blocks (FXD), randomly intermingled within blocks (RND), or randomly intermingled but auditorily cued at 950 ms pre-TMS (RNDC). MEPs showed significant effects of ITI, Pattern, and their interaction, being smaller at 2 s than at longer ITIs in FXD but not in RND and RNDC. However, RNDC produced marked MEP suppression across all ITI. SICI showed a Pattern × ITI interaction but no main effect of Pattern or between-pattern differences per ITI, whereas LICI showed complex Pattern × ITI interaction and both main effects. Notably, ITI affected SICI and LICI oppositely. While GABA-A-receptor mediated inhibition (SICI) showed an ITI dependency compatible with the ITI effects on single-pulse MEPs, it was unlikely mediating the predictability effects but may rather relate to an independent ITI driven suppression. In contrast, GABA-B-receptor mediated inhibition (LICI), was clearly modulated by predictability, but incompatible with the observed ITI effects on MEP amplitude. In summary, the effect of ITI duration on MEP amplitude seems to be largely explained by anticipatory suppression of corticospinal excitability based on the predictability of TMS pulses as derived from their temporal context, being easier for shorter and more regular ITIs and maximal for explicit cues immediately preceding the TMS pulse. SICI and LICI are modulated by ITI as well, with the latter affected by predictability, but neither can explain the observed predictability effects on single-pulse MEP amplitudes.

## 1. Introduction

The effects of transcranial magnetic stimulation (TMS) can be characterized at different spatial levels of neural organization, spanning from local brain circuits (Lazzaro et al., 2008; Massimini et al., 2026) to large-scale brain networks (Fox et al., 2012; Siebner et al., 2009). TMS over the primary motor cortex (M1) evokes motor-evoked potentials (MEPs) via the corticospinal tract, that are measured with electromyography (EMG) and serve as a widely used index of corticospinal excitability (Barker et al., 1985; Hallett, 2007). The MEP amplitude reflects the summed electrical muscle potential of activated muscle units, and despite excitability fluctuations at the spinal level, its amplitude is roughly proportional to the number of corticospinal output neurons in M1 that produce action potentials in response to TMS (Siebner et al., 2022). Beside a considerable amount of yet unexplained spontaneous fluctuations in trial-by-trial MEP amplitude, several physiological and methodological factors have been identified that modulate MEP amplitude. These include for example oscillatory cortical states (Bergmann et al., 2019; Hassan et al., 2025; Thut et al., 2017; Zrenner et al., 2018) and spinal motoneuron excitability fluctuations (Rösler et al., 2008), TMS pulse characteristics, such as intensity (Pellegrini et al., 2018), current direction (Mills et al., 1992), and pulse waveform (Davila-Pérez et al., 2018), and the input from intra- and inter-cortical circuits, which have been extensively studied using paired-pulse single- and dual-coil TMS protocols, in which a conditioning pulse via the same coil over M1 or a different coil over a connected brain region is used to facilitate or inhibit the MEP amplitude of the test pulse depending on its intensity and inter-pulse interval (Cash & Ziemann, 2021; Di Lazzaro & Ziemann, 2013). Importantly, via such large-scale networks, MEP amplitudes are also under top-down control of higher order processes, being increased during action imagery (Facchini et al., 2002; Fadiga et al., 1998), preparation (Leocani et al., 2000), execution (Hess et al., 1987), and observation (Fadiga et al., 1995), and suppressed during response inhibition in No-Go trials (Leocani et al., 2000) or stop-signal tasks (Badry et al., 2009).

Also, the inter-trial interval (ITI) between two subsequent single-pulse TMS-evoked MEPs has been recognized as a critical methodological factor in TMS research, with methodological reviews and expert consensus studies emphasizing the importance of carefully controlling and explicitly reporting its implementation due to its potential influence on the reliability of TMS outcomes (Chipchase et al., 2012; Groppa et al., 2012). Despite this strong consensus, the question of how ITI contributes to MEP variability remains comparatively underexplored relative to other stimulation parameters.

Previous studies have reported mixed effects of ITI on MEP amplitude. In resting muscle, longer intervals have often been associated with larger and less variable MEPs (Hassanzahraee et al., 2019; Julkunen et al., 2012; Matilainen et al., 2022; Vaseghi et al., 2015), whereas other studies found no differences across jittered or separately blocked intervals between 4 and 12 s, including for SICI and intracortical facilitation (Capozio et al., 2021; De Albuquerque et al., 2024; Pantovic et al., 2023). This heterogeneity suggests that ITI duration may interact with the temporal structure in which pulses are delivered.

One potential explanation is that blocked, non-jittered stimulation makes pulse timing predictable, particularly at short regular intervals, thereby engaging rhythm-based prediction and anticipatory suppression (Breska & Deouell, 2017). Consistent with this account, temporally predictable TMS elicits smaller MEPs at rest than unexpected stimulation (Capozio et al., 2021; Sriutaisuk & Franz, 2025; Takei et al., 2005; Tran et al., 2021). Because each TMS pulse also provides salient auditory and somatosensory input (Nikouline et al., 1999), temporal context may generate implicit expectations even without an explicit cue. Previous ITI effects may therefore reflect physical pulse spacing, temporal predictability, or both.

In this study, we sought to replicate previous findings and to disentangle the effects of ITI and temporal expectation and predictability on corticospinal excitability, to determine whether ITI-related differences in MEP amplitude are driven (i) by some sort of short-lived basic neurophysiological after-effect induced in local M1 circuits by the previous TMS pulse and decaying over the time course of approximately 10 s, (ii) by higher-order processes of expectancy and preparation based on temporal predictability, or (iii) by their interaction. We further examined whether these effects may be related to mechanisms of intracortical GABAergic inhibition, specifically SICI as a marker of GABA-A-receptor-mediated inhibition and long-interval intracortical inhibition (LICI) as a marker of GABA-B-receptor–mediated inhibition.

We hypothesized that if predictability is the main driver of ITI effects on MEP amplitude, MEP amplitudes would be smaller for short than for long ITI if presented in blocks without jitter and thus with high predictability for short and lower predictability for longer ITIS. We expected this prediction-mediated effect of ITI to disappear when all ITI were presented within the same block randomly in an intermingled fashion, and thus with generally low predictability, with only potential direct ITI effects (i.e., time since last TMS pulse) remaining. We further expected the reduction of MEP amplitude for short ITIs to reappear when introducing a temporal cue in the intermingled condition, thus restoring high predictability. We did not formulate specific a priori hypotheses regarding the contribution of GABA-A- and GABA-B-mediated inhibitory mechanisms to these effects and therefore investigated these associations in an exploratory manner, but we assumed those effects to be of particular importance in case of any immediate effects of ITI being preserved independent of predictability which would thus rather rely on local intracortical mechanism of inhibition

## 2. Material and Methods

### 2.1. Participants

26 right-handed healthy participants (mean age 25.5 ± 3.5 SD,16 female) participated in the study. Prior to participation, all individuals were screened by a study physician for contraindications to TMS as well as a history of neurological or psychiatric disorders. All measurements were conducted at the Neurostimulation Laboratory of the Neuroimaging Center (NIC) at the University Medical Center Mainz. The study was approved by the ethics committee of the state of Rhineland-Palatinate (No. 2021-15743) and was conducted in accordance with the Declaration of Helsinki. Written informed consent was obtained from all participants prior to study enrollment.

### 2.2. Experimental Design

In a single session, participants completed three experimental blocks (RND, FXD, RNDC), each comprising single-pulse (SP) TMS trials as well as two types of paired-pulse (PP) TMS trials, with paired-pulse conditions using ISIs of 2 ms and 70% RMT for the conditioning pulse (for SICI) and 100 ms and 120% RMT for the conditioning pulse (for LICI) and ITIs of 2, 5, and 10 s. After normalization to the respective single-pulse from the same block and ITI condition, the paired-pulse measures were referred to as SICI and LICI, respectively. Each block thus included all combinations of the three protocols (SP, SICI, LICI) and three ITIs (2, 5, 10 s), with 25 trials for each of the resulting 9 conditions (i.e., protocol–ITI combination) and 225 trials per block.

The order of presenting protocols and ITIs differed for the three experimental blocks according to three predefined expectancy patterns:

1. Randomized pattern (RND; unpredictable): Trials were presented in a randomized order across conditions using MATLAB’s *randperm* function, with no predefined sequence structure.
2. Fixed pattern (FXD; predictable; or rhythmic predictability): Trials were organized into clusters defined by identical ITIs. Within each ITI cluster (of 2, 5, or 10 s), SP and PP trials were repeatedly presented in a fixed order (SP → SICI → LICI). The order of ITI clusters was randomized across participants, while the within-cluster protocol order was identical for all participants.
3. Randomized-cue pattern (RNDC; highly predictable; or arrhythmic predictability): Trials were randomized across conditions as in RND, but each trial was preceded by a consistent auditory cue 950 ms prior to stimulation.

Cues were generated using a second TMS stimulator (MagPro X100, MagVenture), which delivered single pulses to produce a distinct clicking sound serving as a predictive warning signal. The coil was mounted via a FlexArm (MagVenture) on the same side as the active TMS coil targeting M1 but tilted (90°) away from the scalp to prevent cortical stimulation. Cue pulse intensity was set to 85% of maximum stimulator output (MSO). This intensity was chosen so that the click sound could be clearly heard without causing a startle response.

The order of the three experimental blocks was randomized across participants. Each block lasted approximately 21 minutes and was followed by a 15-minute inter-block break, during which participants left the experimental room and took a walk outside to get some fresh air. These breaks helped to reduce potential confounding effects of drowsiness and allowed the TMS coil to cool down. To monitor levels of wakefulness and fatigue throughout the experiment, participants rated their progressive sleepiness before the start of each block using the Stanford Sleepiness Scale (Hoddes et al., 1973). The entire experiment lasted approximately 3.5 hours.

### 2.3. Experimental Procedures

Participants were seated comfortably in a treatment chair (MagVenture) and were instructed to keep their hand muscles relaxed throughout the experiment while maintaining their gaze on a black fixation cross at the wall in front of them. The head was adjusted to an upright position and firmly stabilized with a vacuum pillow. TMS was administered to the hand area of the left primary motor cortex (M1) using a 75 mm figure-of-eight coil (Cool-B-65-CO) connected to a MagPro X100 TMS stimulator with MagOption (MagVenture, Denmark). In 17 participants, the coil was mounted on the TMS-Cobot (Axilum Robotics, France), whereas in 9 participants it was attached to the FlexArm (MagVenture, Denmark) due to technical maintenance of the Cobot. Biphasic pulses were delivered across all trials and a posterior-anterior current was induced in the brain tissue for the second half-wave.

The optimal coil position producing consistent MEPs in the right first dorsal interosseous (FDI) was identified as the motor hotspot using MR-template-based frameless stereotactic neuronavigation (TMS-Navigator, Localite, Germany) and saved for subsequent stimulation. Following hotspot identification, resting motor threshold (RMT) was estimated using the BEST toolbox (Hassan et al., 2022), which implements an automated closed-loop adaptive threshold-hunting (staircase) procedure (Awiszus, 2003). RMT was estimated separately for different ITIs to account for potential variability in RMT estimates across ITI conditions. The final stimulation intensity was defined as the mean of the three RMT estimates for each participant (49.60 ± 7.03 %MSO).

Single pulses were delivered at suprathreshold intensity of 120% RMT. To probe SICI, two pulses separated by an ISI of 2 ms were delivered. The intensity of the subthreshold conditioning stimulus (CS) was set to 70% RMT and the intensity of the suprathreshold test stimulus (TS) was fixed to 120% RMT. We chose a lower intensity for the conditioning pulse, rather than the more commonly used 70% RMT, to prevent ceiling effects in inhibition (Tugin et al., 2021; Vucic et al., 2009). For the LICI protocol, both the CS and TS were presented at suprathreshold intensity with 120% RMT, separated by an ISI of 100 ms (Cash & Ziemann, 2021; Fatih et al., 2021).

### 2.4. Data recording

Surface EMG was recorded from the right FDI, abductor pollicis brevis (APB), and abductor digiti minimi (ADM) muscles. The skin was thoroughly cleaned with alcohol, and disposable surface electrodes were placed in a bipolar belly–tendon montage, with the active electrode positioned over the respective muscle and the reference electrode placed on the bone at the first joint of the corresponding finger. A ground electrode was placed just below the elbow. EMG signals were digitized in DC mode at a 5 kHz sampling rate with a 1250 Hz anti-aliasing low-pass filter, using a TMS-compatible 24-bit amplifier (NeurOne Tesla with Digital-Out Option, Bittium, Finland) powered by a 7.2 V battery.

### 2.5. Data analysis

EMG data were analyzed offline using MATLAB (R2023b) and the FieldTrip toolbox (Oostenveld et al., 2011). Continuous EMG signals from the FDI muscle were first high-pass filtered at 0.5 Hz using a first-order Butterworth filter to remove slow baseline drifts. This filter setting was chosen to avoid potential amplitude distortions, as higher cut-off frequencies (> 1 Hz) or higher filter orders may introduce signal distortions in transient, non-periodic signals such as the MEP waveform (Zschorlich et al., 2021). Continuous EMG signals were then epoched around each TMS pulse using a window extending from 950 ms before to 950 ms after stimulation, providing sufficient temporal padding for subsequent frequency-domain filtering. Powerline interference was removed on a trial-by-trial basis using a 50 Hz discrete Fourier transform (DFT) filter, with signal amplitudes at the line-noise frequency replaced by interpolation from neighboring frequency bins. This procedure prevented spectral discontinuities and minimized distortion of the EMG signal in the vicinity of the line-noise frequency (Leske & Dalal, 2019).

Baseline EMG activity was extracted from a -100 to -5 ms pre-stimulus window. Pre-stimulus EMG signals were smoothed using a five-sample moving-average window to reduce transient spike-like artifacts while preserving the underlying EMG envelope. Trials were excluded if pre-stimulus EMG peak-to-peak amplitude exceeded 50 µV, indicating involuntary muscle activation (Tran et al., 2020, 2021). These criteria ensured that MEP measurements were not confounded by pre-activation of the target muscle.

EMG data were then re-epoched from −50 to +200 ms relative to the TMS pulse for MEP analysis. To correct for DC offset and baseline activity, the signal was demeaned using the mean prestimulus EMG within a −50 to −5 ms window. MEP amplitudes were quantified on a single-trial basis as the peak-to-peak EMG amplitude, calculated as the absolute difference between the minimum and maximum EMG values within a 15 to 50 ms post-stimulus window following the test pulse. For LICI trials, TS MEPs were extracted from the same window shifted by 100 ms to align with the latency of the conditioned test response (Rossini et al., 2015).

Raw MEP amplitudes were log-transformed to reduce positive skewness and stabilize variance. Consistent with conventional paired-pulse TMS methodology, conditioned MEPs evoked by the test pulse of the PP TMS trials were normalized to the corresponding SP MEPs from the same condition (i.e. pattern x ITI combination) (Rossini et al., 2015). Specifically, raw SICI and LICI MEP amplitudes were expressed as a ratio of the corresponding average SP MEP amplitude, whereas log-transformed SICI and LICI MEPs were normalized by subtracting the corresponding average log-transformed single-pulse MEP value. This trial-level normalization approach preserved trial-by-trial variability while accounting for condition-specific differences in baseline corticospinal excitability (Sanger et al., 2001).

For statistical modeling, log-transformed values were used, whereas ratio-normalized values were used for data visualization to facilitate interpretability in the original measurement scale (Bonnesen et al., 2022). All the trials were visually inspected for quality control prior to statistical modeling to ensure signal quality and absence of artifacts.

### 2.6. Statistical Analysis

Statistical analyses were conducted in RStudio Version 2026.4.0.526 (Posit team, 2024), running on R Version 4.4.2 (R Core Team, 2024) using linear mixed-effects models (LMMs) to account for the repeated-measures design and inter-individual variability in MEP amplitudes. Separate models were fitted for each protocol (i.e., SP, SICI, LICI) to obtain protocol-specific effects without imposing cross-protocol interactions. For the SP protocol, the dependent variable was single-trial log-transformed MEP amplitude. For paired-pulse protocols (SICI and LICI), trial-wise normalized MEP amplitudes were calculated by dividing each single-trial test stimulus (TS) MEP amplitude by the corresponding condition-specific mean SP MEP amplitude. These normalized values were then log-transformed and used as the dependent variable in the respective LMMs.

Within each protocol, fixed effects comprised Pattern (three levels: RND, FXD, RNDC), ITI (three levels: 2 s, 5 s, 10 s), and their interaction (Pattern × ITI), forming a 3 × 3 within-subject factorial design. Cumulative session time, computed as the running sum of preceding ITIs, was also included as a nuisance covariate to account for gradual time-dependent changes in corticospinal excitability across the session (e.g., fatigue, habituation, or fluctuations in arousal) and to index experimental progression. The variable was z-standardized prior to modeling to facilitate coefficient interpretation and improve model convergence.

LMMs were fitted using lme4 (Bates et al., 2015) and lmerTest (Kuznetsova et al., 2017). To avoid inflated false positives, and following recommendations for confirmatory mixed-effects modeling (Barr et al., 2013), we initially specified a model with a random intercept for Subject and random slopes for Pattern and ITI, assuming that the effects of these factors may differ across participants.

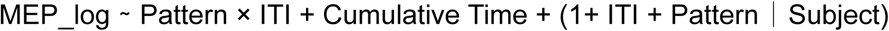

However, this specification resulted in boundary (singular) fits. The estimated variance for the random slope of ITI was close to zero, suggesting limited between-subject variability in the effect of ITI and possible overparameterization of the random-effects structure. We therefore removed the ITI slope while retaining the random slope for Pattern, yielding the final model:

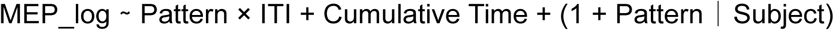

In line with current recommendations for mixed-model analyses of factorial designs, Pattern and ITI were coded using sum-to-zero (effects) contrasts, such that their coefficients represent deviations from the (unweighted) grand mean rather than differences from an arbitrary reference level (Meteyard & Davies, 2020; Schad et al., 2020). Models were estimated by restricted maximum likelihood (REML). Fixed-effect tests for Pattern, ITI, their interaction, and Cumulative Time were obtained using Type III ANOVA with Satterthwaite’s approximation to estimate degrees of freedom, as implemented in lmerTest (Kuznetsova et al., 2017).

Model diagnostics and fit indices, including AIC, marginal and conditional R², intraclass correlation coefficients, and residual error, were computed using the performance (Lüdecke et al., 2021) package. These metrics were used to assess model fit and parsimony, and to confirm that the final specification provided an adequate description of the observed data without indications of misspecification or unnecessary complexity. To further evaluate generalizability beyond the estimation sample, we additionally conducted grouped k-fold cross-validation (k = 10) using the caret package (Kuhn, 2008), with all trials from a given participant assigned to the same fold. Cross-validated root mean squared error (RMSE) and mean absolute error (MAE) were used as out-of-sample performance metrics. The consistency of these values across folds indicated stable predictive performance and suggested that model estimates were not unduly influenced by any particular subset of participants or trials.

Where omnibus tests indicated meaningful main effects or interactions involving Pattern and/or ITI, we conducted planned post-hoc comparisons of estimated marginal means using the emmeans package (Lenth, 2025). Specifically, we obtained (i) all pairwise comparisons among ITI levels averaged over Pattern, (ii) all pairwise comparisons among Pattern levels averaged over ITI, and (iii) simple-effect contrasts of ITI within each Pattern level and Pattern within each ITI level to unpack the Pattern × ITI interaction. All sets of pairwise comparisons were adjusted for multiple testing using Tukey’s method (Benjamini & Braun, 2002), with the Tukey–Kramer (Kramer, 1956) extension applied to account for unequal group sizes, thereby controlling the familywise error rate within each family of comparisons.

To investigate whether ITI influences RMT, a one-way repeated-measures ANOVA was performed with ITI levels as the within-subject factor. The model was fitted in MATLAB (R2023b) using the *fitrm* function, and the significance of the ITI effect was assessed using the *ranova* function. Following a significant main effect, post hoc pairwise comparisons were conducted using the *multcompare* function with Tukey-Kramer (Kramer, 1956) correction for multiple comparisons. Statistical significance was defined as p < .05 (two-sided) for adjusted p-values.

## 3. Results

Overall, 96.5% of recorded trials were retained for analysis, while the remaining 3.5% were excluded due to technical issues (.96%) or pre-stimulus EMG activity (2.54%).

### 3.1. Effect of ITI on RMT

Mean RMT values (mean ± SD, %MSO) were 51.62 ± 7.88, 49.35 ± 7.34, and 47.85 ± 6.93 at ITIs of 2 s, 5 s, and 10 s, respectively. A one-way repeated-measures ANOVA revealed a significant effect of ITI on RMT, F(2, 50) = 11.76, p < .001. Post hoc pairwise comparisons with correction for multiple comparisons showed that RMT was significantly higher at the 2 s ITI than at both the 5 s ITI (p = .005) and the 10 s ITI (p = .001). No significant difference was observed between the 5 s and 10 s ITIs (p = .130). These findings indicate that shorter ITIs are associated with higher RMT estimates, whereas extending the ITI from 5 s to 10 s does not produce a further significant change in RMT.

### 3.2. SP

Linear mixed-effects modeling revealed significant main effects of ITI, F(2, 5617.01) = 28.57, p < .001, and Pattern, F(2, 24.97) = 16.14, p < .001, as well as a significant ITI × Pattern interaction, F(4, 5616.91) = 37.52, p < .001, indicating that the effect of ITI on corticospinal excitability depended on stimulation predictability.

To follow up on the main effect of ITI (**Figure 1A**), pairwise comparisons of estimated marginal means (averaged across Pattern levels) showed that MEP amplitudes were significantly lower at the 2 s ITI than at the 5 s (Estimate = −.10, 95% CI [−.13, −.07], p < .001) and the 10 s ITIs (Estimate = −.07, 95% CI [−.11, −.04], p < .001), indicating reduced corticospinal excitability at the 2 s ITI. No significant difference was observed between the 5 s and 10 s ITIs (p = .100).

**Figure 1:**
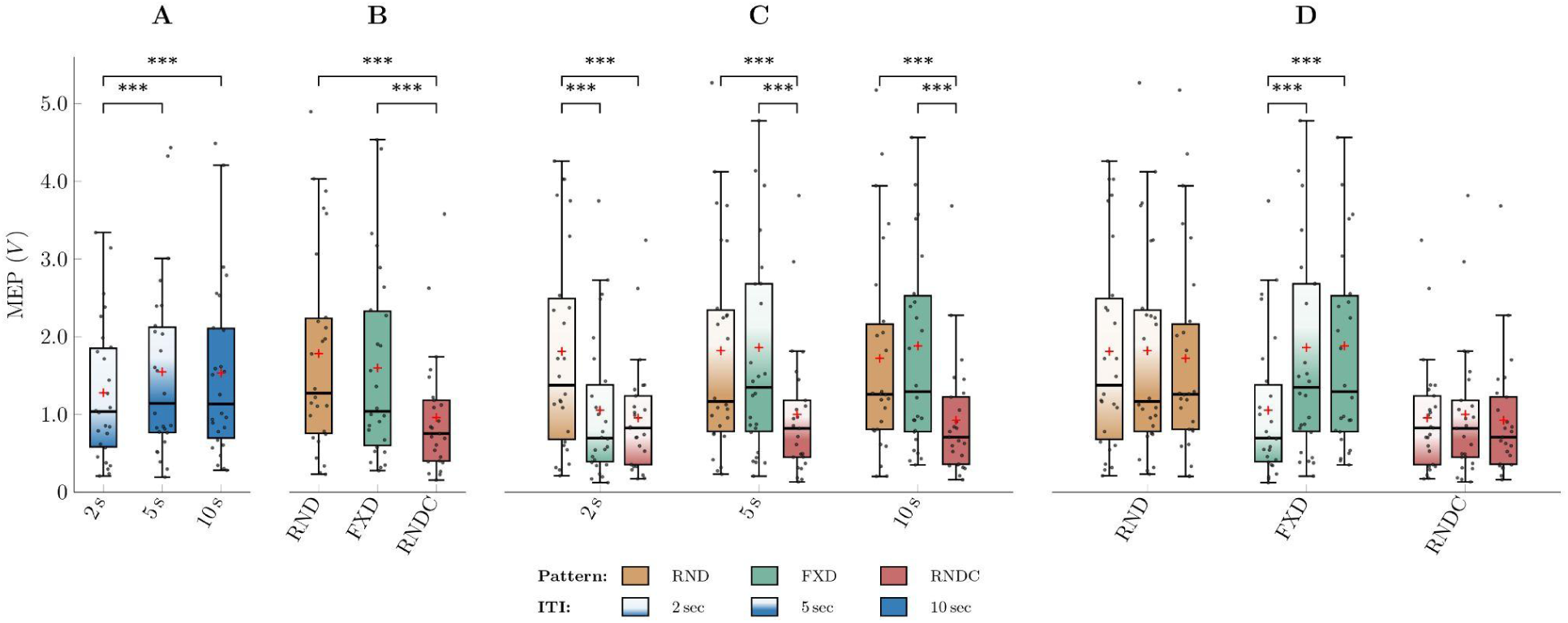
Effects of temporal predictability and ITI on single-pulse MEPs. MEP amplitudes are shown across stimulation patterns and ITIs. **(A)** MEPs across ITIs, collapsed across stimulation patterns. **(B)** MEPs across stimulation patterns, collapsed across ITIs. **(C)** MEPs across stimulation patterns separately for each ITI. **(D)** MEPs across ITIs separately for each stimulation pattern. Across the figure, progressively darker shading within each stimulation pattern denotes increasing ITI (2 s, 5 s, and 10 s, respectively). In each panel, boxplots show the distribution of subject-level MEP values. The box represents the interquartile range (IQR), the horizontal line indicates the median, and the whiskers extend to the most extreme values within 1.5 × IQR. Black dots represent individual participants, and the plus sign (+) indicates the mean normalized MEP at each ITI or stimulation-pattern level. Statistical significance is indicated as follows: ∗∗∗ = p < .001, ∗∗ = p < .01, ∗ = p < .05.

To further examine the main effect of Pattern (**Figure 1B**), pairwise comparisons of estimated marginal means (averaged across ITI levels) showed that MEP amplitudes were significantly lower in the highly predictable RNDC than in both the predictable FXD (Estimate = .34, 95% CI [.17, .51], p < .001) and the unpredictable RND patterns (Estimate = .42, 95% CI [.23, .60], p < .001), indicating reduced corticospinal excitability in the RNDC condition. No significant difference was observed between the FXD and RND conditions (p = .120).

Pairwise comparisons of ITI effects within each Pattern level (**Figure 1D**) showed that corticospinal excitability did not vary significantly across ITIs in either the low-predictability RND condition (all p ≥ .251) or the highly predictable RNDC condition (all p ≥ .156, although the 2 s versus 10 s comparison reached nominal significance, p = .034). In contrast, within the FXD condition, corticospinal excitability was reduced at the 2 s ITI relative to both the 5 s (Estimate = −.29, 95% CI [−.35, −.24], p < .001) and 10 s ITIs (Estimate = −.29, 95% CI [−.35, −.23], p < .001), with no difference between the 5 s and 10 s ITIs (p = .992).

Pairwise comparisons of Pattern effects within each ITI level (**Figure 1C**) showed a similar pattern at the 5 s and 10 s ITIs. At both levels, the highly predictable RNDC condition elicited significantly lower MEP than both FXD and RND (5 s ITI: FXD vs. RNDC: Estimate = .43, 95% CI [.25, .61], p < .001; RND vs. RNDC: Estimate = .43, 95% CI [.24, .62], p < .001; 10 s ITI: FXD vs. RNDC: Estimate = .47, 95% CI [.29, .65], p < .001; RND vs. RNDC: Estimate = .43, 95% CI [.25, .62], p < .001). In contrast, at the 2 s ITI, MEP was higher in the RND condition compared with both FXD (FXD vs. RND: Estimate = −.26, 95% CI [−.37, −.16], p < .001) and RNDC (RND vs. RNDC: Estimate = .38, 95% CI [.20, .57], p < .001).

### 3.3. SICI

Linear mixed-effects modeling revealed a significant main effect of ITI, F(2, 5547.99) = 7.18, p = .001, indicating that MEP amplitude varied as a function of stimulation interval. In contrast, neither the main effect of Pattern, F(2, 25.16) = 0.74, p = .486, nor the overall differences in expectancy across patterns (all p ≥ .449, **Figure 2B**) reached significance. Although the ITI × Pattern interaction was statistically significant, F(4, 5547.86) = 2.57, p = .036, the effect was modest and pairwise comparisons did not reveal a consistent modulation across pattern conditions.

**Figure 2:**
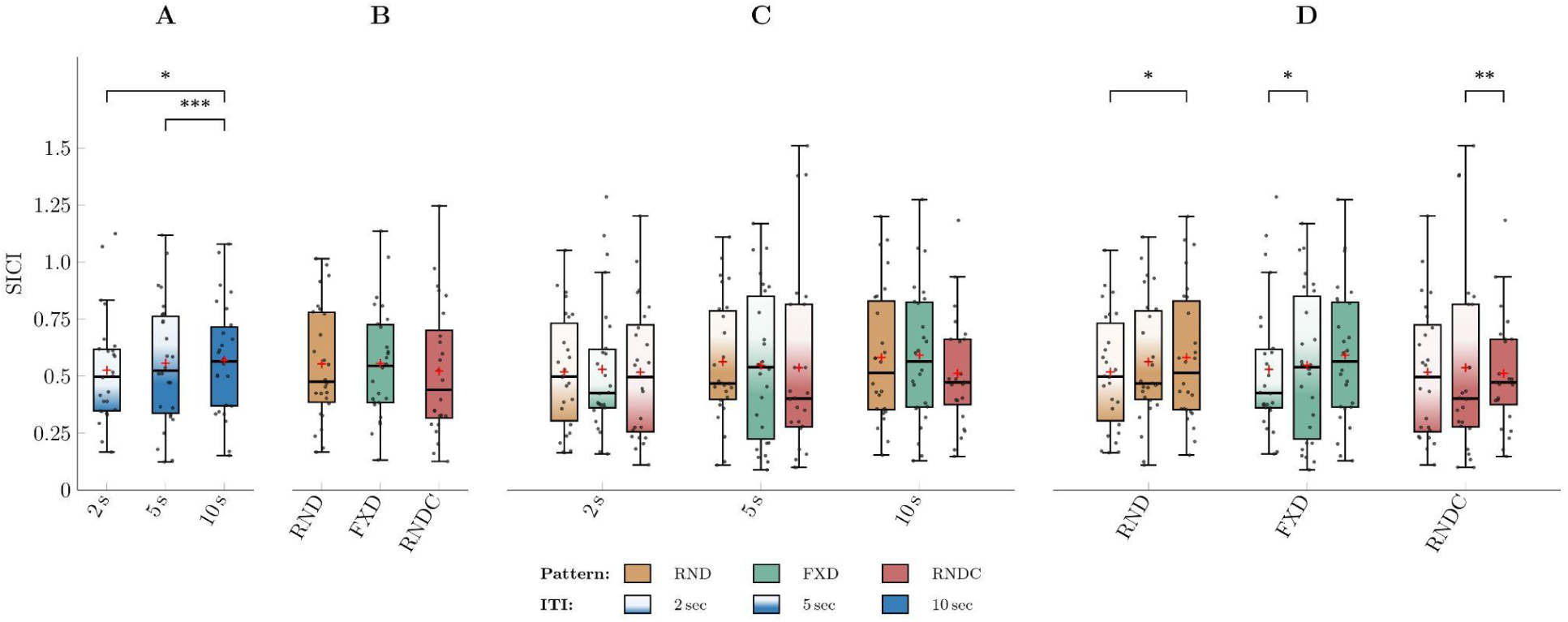
Effects of temporal predictability and ITI on SICI. SICI is shown as the ratio of the conditioned MEP to the corresponding unconditioned MEP obtained within the same stimulation pattern and ITI condition. Lower normalized values indicate stronger SICI, whereas higher values indicate weaker inhibition. **(A)** SICI MEPs across ITIs, collapsed across stimulation patterns. **(B)** SICI MEPs across stimulation patterns, collapsed across ITIs. **(C)** SICI MEPs across stimulation patterns separately for each ITI. **(D)** SICI MEPs across ITIs separately for each stimulation pattern. Across the figure, progressively darker shading within each stimulation pattern denotes increasing ITI (2 s, 5 s, and 10 s, respectively). In each panel, boxplots show the distribution of subject-level MEP values. The box represents the interquartile range (IQR), the horizontal line indicates the median, and the whiskers extend to the most extreme values within 1.5 × IQR. Black dots represent individual participants, and the plus sign (+) indicates the mean normalized MEP at each ITI or stimulation-pattern level. Statistical significance is indicated as follows: ∗∗∗ = p < .001, ∗∗ = p < .01, ∗ = p < .05.

Across ITIs (**Figure 2A**), conditioned MEP amplitudes were significantly larger at the 10 s ITI than at both the 2 s (Estimate = −.04, 95% CI [−.08, −.00], p = .026) and 5 s ITIs (Estimate = −.06, 95% CI [−.09, −.02], p = .001), indicating reduced SICI at the longest ITI. The remaining pairwise comparison was not significant (p = .517).

Follow-up analyses revealed significant ITI-related differences across stimulation patterns, although the pattern of pairwise effects varied (**Figure 2D**). In RND, conditioned MEP amplitudes were higher at 10 s than at 2 s (Estimate = −.07, 95% CI [−.13, .00], p = .029), whereas in RNDC, amplitudes were higher at 10 s than at 5 s (Estimate = −.09, 95% CI [−.15, −.03], p = .003). In FXD, the only significant difference was between 2 s and 5 s (Estimate = .06, 95% CI [.00, .12], p = .047). All other within-pattern pairwise comparisons were not significant (all p ≥ .069).

Despite these isolated pairwise effects, direct comparisons between stimulation patterns at each ITI (**Figure 2C**) revealed no significant differences (all p ≥ .203).

Overall, SICI modulation varied primarily as a function of ITI, whereas no main effect of stimulation Pattern was observed. Although the ITI × Pattern interaction was statistically significant, the pattern-specific pairwise effects were sparse and inconsistent, with different ITI contrasts reaching significance across patterns and no significant between-pattern differences at any ITI. Thus, the interaction did not reveal a systematic modulation of SICI by pattern predictability.

### 3.4. LICI

Linear mixed-effects modeling revealed significant main effects of ITI, F(2, 5158.53) = 22.80, p < .001, and Pattern, F(2, 23.96) = 15.47, p < .001, as well as a significant ITI × Pattern interaction, F(4, 5158.00) = 34.48, p < .001, indicating that the effect of ITI on LICI varied across stimulation patterns.

Pairwise comparisons of ITIs, collapsed across Pattern levels (**Figure 3A**), showed that LICI differed between 2 s and 5 s (Estimate = .09, 95% CI [.06, .12], p < .001) and between 2 s and 10 s (Estimate = .06, 95% CI [.03, .10], p < .001), whereas the 5 s and 10 s ITIs did not differ (p = .128). Thus, LICI was higher at 5,s and 10,s than at 2,s, indicating reduced inhibition at the longer ITIs.

**Figure 3:**
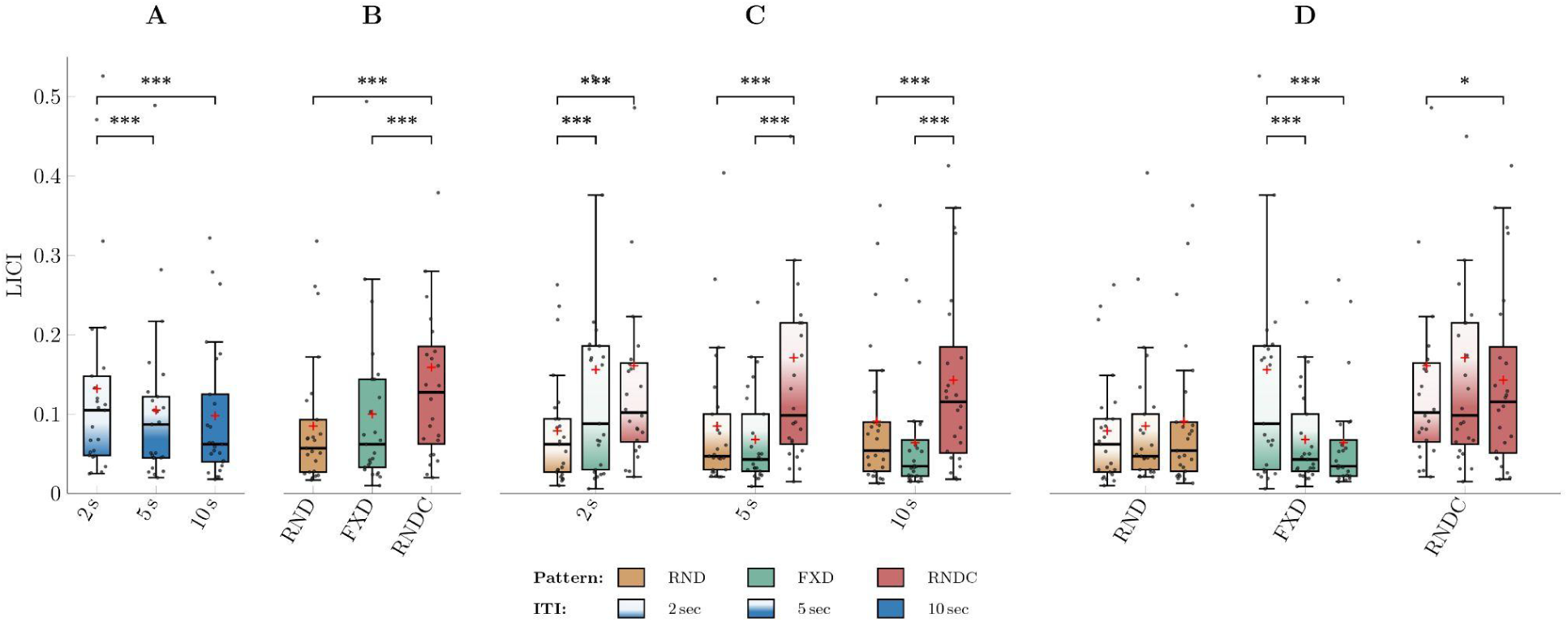
Effects of temporal predictability and ITI on LICI. LICI is shown as the ratio of the test-stimulus MEP, elicited 100 ms after the conditioning stimulus, to the corresponding unconditioned MEP obtained within the same stimulation pattern and ITI condition. Lower normalized values indicate stronger LICI, whereas higher values indicate weaker inhibition. **(A)** LICI MEPs across ITIs, collapsed across stimulation patterns. **(B)** LICI MEPs across stimulation patterns, collapsed across ITIs. **(C)** LICI MEPs across stimulation patterns separately for each ITI. **(D)** Normalized LICI MEPs across ITIs separately for each stimulation pattern. Across the figure, progressively darker shading within each stimulation pattern denotes increasing ITI (2 s, 5 s, and 10 s, respectively). In each panel, boxplots show the distribution of subject-level MEP values. The box represents the interquartile range (IQR), the horizontal line indicates the median, and the whiskers extend to the most extreme values within 1.5 × IQR. Black dots represent individual participants, and the plus sign (+) indicates the mean normalized MEP at each ITI or stimulation-pattern level. Statistical significance is indicated as follows: ∗∗∗ = p < .001, ∗∗ = p < .01, ∗ = p < .05.

Pairwise comparisons of stimulation Patterns, collapsed across ITI levels (**Figure 3B**), showed that LICI was lower in RNDC than in both FXD (Estimate = −.33, 95% CI [−.51, −.14], p < .001) and RND (Estimate = −.39, 95% CI [−.56, −.21], p < .001), whereas FXD and RND did not differ (p = .307). Thus, RNDC showed reduced GABA-B-mediated intracortical inhibition, while FXD and RND showed comparable levels of inhibition.

Pairwise comparisons of ITIs within each stimulation Pattern (**Figure 3D**) showed that ITI-related changes in LICI varied across Patterns. In RND, no ITI comparisons were significant (all p ≥ .382). In FXD, LICI was higher at both 5 s (Estimate = .28, 95% CI [.22, .33], p < .001) and 10 s (Estimate = .28, 95% CI [.22, .34], p < .001) than at 2 s, with no difference between 5 s and 10 s (p = .974). In RNDC, LICI was lower at 10 s than at 2 s (Estimate = −.06, 95% CI [−.12, −.00], p = .028), with no other significant differences (all p ≥ = .053). Thus, ITI-related modulation of LICI differed between FXD and RNDC, with FXD showing increased inhibition at longer ITIs, whereas RNDC tended toward reduced inhibition at longer ITIs.

Pairwise comparisons between stimulation Patterns at each ITI (**Figure 3C**) showed that at both 5 s and 10 s, RNDC had lower LICI than both FXD (5 s: Estimate = −.40, 95% CI [−.59, −.21]; 10 s: Estimate = −.46, 95% CI [−.65, −.27]; both p < .001) and RND (5 s: Estimate = −.38, 95% CI [−.56, −.20]; 10 s: Estimate = −.41, 95% CI [−.58, −.23]; both p < .001), indicating reduced GABA-B-mediated inhibition in RNDC. At 2 s, RND had higher LICI than both FXD (FXD vs. RND: Estimate = .25, 95% CI [.15, .36], p < .001) and RNDC (Estimate = −.37, 95% CI [−.55, −.19], p < .001), indicating greater GABA-B-mediated inhibition in RND. All other between-pattern comparisons were not significant (all p ≥ .275).

### 3.5. Effect of Cumulative Experimental Time on MEP Amplitude

Cumulative experimental time was significantly associated with log-transformed MEP amplitude in the SP–LICI CS model (F(1, 11052.23) = 50.88, β = .029, p < .001), indicating that a one standard deviation increase in cumulative experimental time was associated with an approximately 2.9% increase in MEP amplitude. Similarly, cumulative time was positively associated with MEP amplitude for SP (F(1, 5620.31) = 23.90, β = .029, p < .001) and SICI (F(1, 5627.49) = 29.22, β = .037, p < .001), whereas a negative association was observed for LICI (F(1, 5379.97) = 22.17, β = −.029, p < .001). These findings indicate that MEP amplitudes exhibited condition-dependent temporal changes across the experimental session. Inclusion of cumulative time improved model fit across analyses, as reflected by lower AIC values, demonstrating that temporal variation contributed to explaining MEP variability. However, the magnitude of this effect was small, and inclusion of the covariate resulted in only minimal changes in the estimated fixed effects, indicating that the main experimental effects remained robust to temporal trends.

## 4. Discussion

### 4.1. Summary of key findings

The main finding was that the effect of ITI on single-pulse MEP amplitude depended strongly on temporal context. In RND, MEPs did not differ across the 2, 5, and 10 s ITIs. In FXD, MEPs were smaller at 2 s than at 5 or 10 s, whereas RNDC produced marked suppression across ITIs, particularly relative to both uncued conditions at 5 and 10 s. RMT was likewise higher at 2 s than at the two longer intervals. SICI ratios varied mainly with ITI, with stronger inhibition at shorter intervals when collapsed across patterns, but showed no systematic effect of predictability. LICI ratios showed pronounced effects of ITI, Pattern, and their interaction, yet their condition-specific profile did not reproduce the single-pulse MEP pattern. Together, these findings indicate that pulse spacing alone does not determine corticospinal output and favor temporal predictability as a major contributor to the observed MEP suppression; neither paired-pulse index, however, provided a sufficient mechanistic explanation.

### 4.2. Effects of temporal context and predictability on MEP amplitudes

Previous single-pulse studies do not support a simple, context-independent dose-response to ITI. In resting muscle, fixed ITIs from 1 to 10 s or from 4 to 10 s produced progressively larger MEPs with increasing interval (Julkunen et al., 2012; Vaseghi et al., 2015), and extending ITIs from 5 s to 10–20 s increased MEP amplitude and reduced variability, with little further gain beyond approximately 15 s (Hassanzahraee et al., 2019). Matilainen et al. (2022) likewise observed smaller MEPs at 2 s than at 5 or 10 s in resting muscle, but no ITI effect during voluntary contraction. In contrast, Capozio et al. (2021) found no amplitude difference between jittered short and long intervals, and separately blocked ITIs of 4, 6, 8, and 10 s did not systematically alter unconditioned MEPs, SICI, or intracortical facilitation in two recent studies (De Albuquerque et al., 2024; Pantovic et al., 2023). Differences in muscle state, interval range, trial order, jitter, and block structure may therefore explain part of the apparent inconsistency. In particular, many studies reporting smaller MEPs at short ITIs used fixed, blocked trains, which confound elapsed time since the previous pulse with rhythmic predictability.

The present manipulation helps separate these factors. The absence of an ITI effect in RND shows that a 2 s interval was not sufficient, by itself, to suppress corticospinal output. The selective reduction at 2 s in FXD instead indicates that rhythmic regularity made the shortest interval particularly effective, while the explicit warning cue in RNDC suppressed MEPs across the full ITI range. This interpretation converges with studies showing reduced MEPs when TMS timing was predictable, whether timing was explicitly signaled, learned from a reliable cue, or self-initiated (Capozio et al., 2021; Sriutaisuk & Franz, 2025; Takei et al., 2005; Tran et al., 2021). Capozio et al. (2021) also found that attenuating the TMS click reduced MEP amplitude, illustrating that auditory input and stimulus expectation can exert distinct, potentially opposing influences. The RNDC effect therefore cannot be assigned exclusively to temporal prediction: the sham-TMS click may additionally have induced auditory orienting, alerting, or sensory priming. Likewise, RND was not devoid of temporal information, because the conditional probability of stimulation increased as the shorter intervals elapsed. The most defensible conclusion is that temporal context and cueing strongly modulate MEP amplitude, with anticipatory suppression favored by short regular intervals and maximized by a precise warning event, while the sensory contribution of that event remains unresolved.

### 4.3. Resting motor threshold

The RMT results show a closely related short-interval profile. RMT was higher at 2 s than at both 5 s and 10 s, with no difference between the two longer intervals, indicating reduced responsiveness during short-interval threshold estimation. Because RMT was estimated separately for each ITI, each estimate was obtained in a temporally regular context resembling FXD. The parallel between higher RMT and smaller suprathreshold MEPs at 2 s in FXD is therefore consistent with the same combination of short spacing and rhythmic predictability affecting both measures. This interpretation also agrees with prior evidence that short intervals can reduce resting MEP amplitude or influence individual threshold estimates (Hassanzahraee et al., 2019; Julkunen et al., 2012; Matilainen et al., 2022). However, RMT and suprathreshold MEP amplitude probe different parts of the input–output function, and the RMT procedure did not independently manipulate interval sequence or cueing. The threshold result therefore supports the practical conclusion that a 2 s ITI can bias excitability estimates in a regular train, but it cannot determine whether spacing, predictability, or their interaction caused the effect.

### 4.4. Potential neurophysiological mechanisms assessed by SICI and LICI

ITI-related effects. Across patterns, SICI ratios were lower at 2 and 5 s than at 10 s, indicating stronger relative inhibition at the shorter intervals according to the ratio definition. This overall direction is compatible with the lower single-pulse MEP amplitude at 2 s and could, in principle, contribute to an ITI-linked suppression. The correspondence was not condition-specific, however: within FXD, SICI differed only between 2 and 5 s and did not reproduce the single-pulse contrast between 2 s and both longer intervals, while different contrasts emerged in RND and RNDC. Earlier work likewise found no systematic SICI differences across separately blocked ITIs of 4–10 s (D Albuquerque et al., 2024), suggesting that any ITI effect on SICI may be most apparent at very short intervals. SICI may therefore index an independent ITI-sensitive GABA-A-repeator mediated process, but the present data do not establish it as the mediator of the single-pulse effect. LICI ratios showed an even more condition-specific course: they were higher at 5 and 10 s than at 2 s in FXD, lower at 10 s than at 2 s in RNDC, and unchanged across ITIs in RND. The absence of a common ITI response across patterns argues against a uniform GABA-B-sensitive mechanism driven only by elapsed time since the preceding trial.

Predictability-related effects. SICI showed neither a main effect of pattern nor any between-pattern difference at an individual ITI, despite the small interaction. Thus, SICI did not seem to underlie a strong ITI-dependency in the FXD condition nor a complete lack thereof in the RND condition, and also did not show a strong suppression of single-pulse MEPs in RNDC. It is thus unlikely to mediate the predictability effect. One possibility is that temporal expectation changes the pre-pulse excitability state without substantially altering the incremental effect of a subthreshold conditioning stimulus delivered only 2 ms before the test pulse; in that sense, the fast GABA-A-sensitive process probed by SICI may be comparatively insensitive to the expectation manipulation. However, this explanation remains speculative. LICI, by contrast, clearly distinguished temporal contexts, but its interaction did not reproduce the simple single-pulse signature of FXD-specific suppression at 2 s and RNDC suppression across ITIs. Notably, LICI differs fundamentally from SICI because its suprathreshold conditioning pulse evokes a corticospinal response and salient auditory and somatosensory input 100 ms before the test pulse. The conditioning pulse could therefore partly reset the pre-pulse state or act as a warning event for the test pulse. Such cueing and sensory consequences could make the LICI response reflect processing of the second pulse as well as GABA-B-sensitive inhibition, thereby decoupling it from the single-pulse pattern.

Measurement-related explanations must also be considered. Conventional amplitude-ratio SICI and LICI are sensitive to the neural populations recruited by the test stimulus and are not receptor-specific readouts of GABA-A or GABA-B activity. For LICI, Sailer et al. (2002) directly showed that inhibition decreased when the unconditioned test MEP was increased to a very large amplitude, whereas inhibition was similar for target MEPs of approximately 0.2 and 1 mV. This does not support a simple account in which smaller test MEPs necessarily produce weaker LICI. Nevertheless, the present ratios were normalized to condition-specific single-pulse MEPs, which themselves changed with pattern and ITI; a ratio can therefore change because of its conditioned numerator, its unconditioned denominator, or non-proportional changes in both. A joint analysis of raw single-pulse and conditioned test-pulse MEPs, together with analysis of the LICI conditioning-pulse MEP as a single-pulse response, would distinguish these possibilities and test whether the conditioning pulse resets the predictive state. Until those analyses and the direction of the reported LICI ratios are reconciled, the conservative conclusion is that SICI and LICI are modulated by temporal context but do not yet identify the inhibitory mechanism responsible for anticipatory single-pulse MEP suppression.

### 4.5. Limitations

Several limitations constrain the interpretation of these findings. First, the RNDC cue introduced an additional auditory event, and no trial-wise expectancy measure was obtained; temporal prediction, alerting, and sensory priming therefore cannot be separated in this condition. Second, the finite set of ITIs created an increasing temporal hazard in RND, so predictability was reduced rather than eliminated. Third, normalization of paired-pulse responses to condition-specific single-pulse MEPs makes SICI and LICI ratios sensitive to changes in both numerator and denominator and limits receptor-level interpretation. Fourth, single-pulse, SICI, and LICI trials followed a fixed order within FXD blocks but were intermingled in the other patterns, leaving protocol order and carry-over as potential contributors to pattern differences. Fifth, the RMT procedure did not orthogonally manipulate ITI duration and temporal predictability. Finally, all conditions were acquired in one session. Although cumulative experimental time was modeled and only minimally changed the principal fixed effects, fatigue, vigilance, and other slow state changes may still have contributed. These limitations motivate replication with counterbalanced protocol order, matched predictive and non-predictive auditory control cues, explicit expectancy ratings, and analyses of raw as well as normalized paired-pulse responses.

### 4.6. Conclusions

ITI-related changes in single-pulse MEP amplitude were not determined by pulse spacing alone but depended strongly on the temporal and sensory context of stimulation. Randomizing ITIs abolished the short ITI suppression observed during fixed stimulation, whereas a precise warning cue produced robust suppression across the full ITI range. Together with the parallel RMT result in a regular temporal context, these findings identify predictability as a major determinant of apparent ITI effects, although the auditory contribution of the cue remains unresolved. SICI showed an ITI-sensitive pattern but did not follow the predictability manipulation, while LICI was strongly context dependent but may have been also shaped by its suprathreshold conditioning pulse and by condition-specific normalization. Neither paired-pulse index therefore provides a sufficient mechanistic explanation for anticipatory MEP suppression. Standardizing and reporting ITI duration, interval sequencing, warning cues, and paired-pulse normalization is essential for the interpretation and reproducibility of TMS measures of corticospinal excitability.

## AUTHOR CONTRIBUTIONS

**Saman Seifpour:** Conceptualization, Methodology, Investigation, Software, Formal Analysis, Data Curation, Writing - original draft, Writing - review & editing, and Visualisation. **Nele Haferkorn:** Conceptualization, Methodology, Investigation, Data Curation, Writing - original draft. **Suhas Vijayakumar:** Investigation, Data Curation, Writing – review & editing. **Til Ole Bergmann:** Conceptualization, Methodology, Supervision, Writing – review & editing.

## ACKNOWLEDGEMENTS

AI (GPT-5.6 Sol and Consensus) was used to help refine the writing style and reduce the word count across several sections of this paper. The authors subsequently reviewed and edited these sections and take full responsibility for their content.

## DATA AVAILABILITY STATEMENT

The source code and raw data will be made publicly available on the Open Science Framework (OSF) upon publication of the manuscript.

